# Suppressors of mRNA decapping defects isolated by experimental evolution ameliorate transcriptome disruption without restoring mRNA decay

**DOI:** 10.1101/2020.06.14.151068

**Authors:** Minseon Kim, Ambro van Hoof

## Abstract

Faithful degradation of mRNAs is a critical step in gene expression, and eukaryotes share a major conserved mRNA decay pathway. In this major pathway, the two rate determining steps in mRNA degradation are the initial gradual removal of the poly(A) tail, followed by removal of the cap structure. Removal of the cap structure is carried out by the decapping enzyme, containing the Dcp2 catalytic subunit. While the mechanism and regulation of mRNA decay is well-understood, the consequences of defects in mRNA degradation are less clear. Dcp2 has been reported as either essential or nonessential. Here we clarify that Dcp2 is essential for continuous growth and use experimental evolution to identify suppressors of this essentiality. We show that null mutations in at least three different are each sufficient to restore viability to a *dcp2*Δ, of which *kap123*Δ and *tl(gag)g*Δ appear the most specific. Unlike previously reported suppressors of decapping defects, these suppressor do not restore decapping or mRNA decay to normal rates, but instead allow survival while only modestly affecting transcriptome homeostasis. These effects are not limited to mRNAs, but extend to ncRNAs including snoRNAs and XUTs. These results provide important new insight into the importance of decapping and resolves previously conflicting publications about the essentiality of *DCP2*.

## INTRODUCTION

Eukaryotes share two major mRNA decay pathways that are both carried out by exonucleolytic digestion. mRNA degradation is initiated by gradual shortening of the poly(A) tail, followed by Xrn1-mediated 5′ to 3′ decay and RNA exosome-mediated 3′ to 5′ decay [1]. Because Xrn1 can only degrade RNAs with a 5′-monophosphate [2, 3], removal of the 5′ cap structure by Dcp2 is required in the 5′ to 3′ decay pathway. Importantly, deleting either *DCP2* or *XRN1* results in stabilization of many yeast mRNAs [4–6]. The stabilization of mRNAs in *dcp2* mutants indicates that yeast Dcp2 is the major decapping enzyme, and the 5′ to 3′ pathway is the major mRNA decay pathway. Other enzymes capable of decapping mRNAs have been described both in yeast and other organisms [7–13] but their role in bulk cytoplasmic mRNA degradation has not been fully defined. Consistent with its importance for mRNA decay, deletion of *XRN1* causes a slow growth defect, while the phenotype of *dcp2*Δ is reported inconsistently among different studies. Some studies have reported that *dcp2*Δ is viable but slow-growing, while others reported that *dcp2*Δ is lethal [5, 14–16]. It has been speculated that this difference between studies is attributable to differences between the strains used [16], but this has not been critically analyzed.

Previously, suppressor screens of budding yeast decapping mutants (*dcp1* or *dcp2* conditional mutants) have identified *EDC1*, *EDC2*, *EDC3*, *SBP1*, and *DCP2* itself as high-copy suppressors [5, 17–19]. In each case, the improved growth caused by suppressors was correlated with improved decapping activity and mRNA degradation, suggesting that the essential function of the Dcp1-Dcp2 decapping enzyme is indeed mRNA decapping. While these studies showed that the major function of Dcp1 and Dcp2 is mRNA decapping, they are limited to high-copy suppressor screen of conditional alleles in the decapping enzyme, which may not have revealed the full functions of Dcp2.

To further understand the function of Dcp2, we sought to identify suppressors of the growth defect of a decapping mutant by a complementary experimental evolution of a *dcp2* null strain, which can be more powerful in identifying smaller effects and double mutants. Surprisingly, we identified genes that have no obvious connection to mRNA degradation. Among the genes we identified, we focused on *KAP123* and *tL(GAG)G* that are recurrently mutated. We showed that a null mutation of each gene is sufficient to suppress the lethality of *dcp2*Δ and that *kap123*Δ and *tl(gag)g*Δ have additive effects. We also show that previously reported viable *dcp1*Δ and *dcp2*Δ strains had undetected mutations in *KAP123*, suggesting that they were mistakenly reported as viable due to the suppressor mutations. Instead, our results suggest that *DCP2* is an essential gene. Interestingly, suppression of the growth defect of *dcp2*Δ is not caused by improved cytoplasmic mRNA decay. Instead, our results indicate that absence of Dcp2 causes a global disturbance of the transcriptome including not only mRNA but also non-coding RNA. We also show that the suppressor mutations we identified have a transcriptome-wide but modest amplitude effect in restoring RNA homeostasis. These results indicate that while Dcp2 is normally essential, the essentiality can be overcome by relatively modest adjustments in the transcriptome.

## RESULTS

### *DCP2* is required for the normal growth of yeast

While *DCP2* is annotated as an essential gene in the Saccharomyces Genome Database (yeastgenome.org), there are other studies showing that *DCP2* is not an essential gene [5, 14–16]. The genome-wide effort to identify essential genes was based on sporulating a heterozygous diploid *DCP2/dcp2*Δ strain and attempting to recover haploid *dcp2*Δ strains. We obtained this same commercially available heterozygous diploid *DCP2/dcp2*Δ strain and performed sporulation and tetrad dissection (Figure 1A). We expected that this would produce viable wild-type and inviable *dcp2*Δ progeny in a 1:1 ratio. However, upon prolonged growth we isolated viable *dcp2*Δ (34%) along with wild-type (51%) and inviable *dcp2*Δ progeny (14%) (Figure 1B). Although we were able to recover viable *dcp2*Δ progeny, these spores formed much smaller colonies.

**Figure 1.**
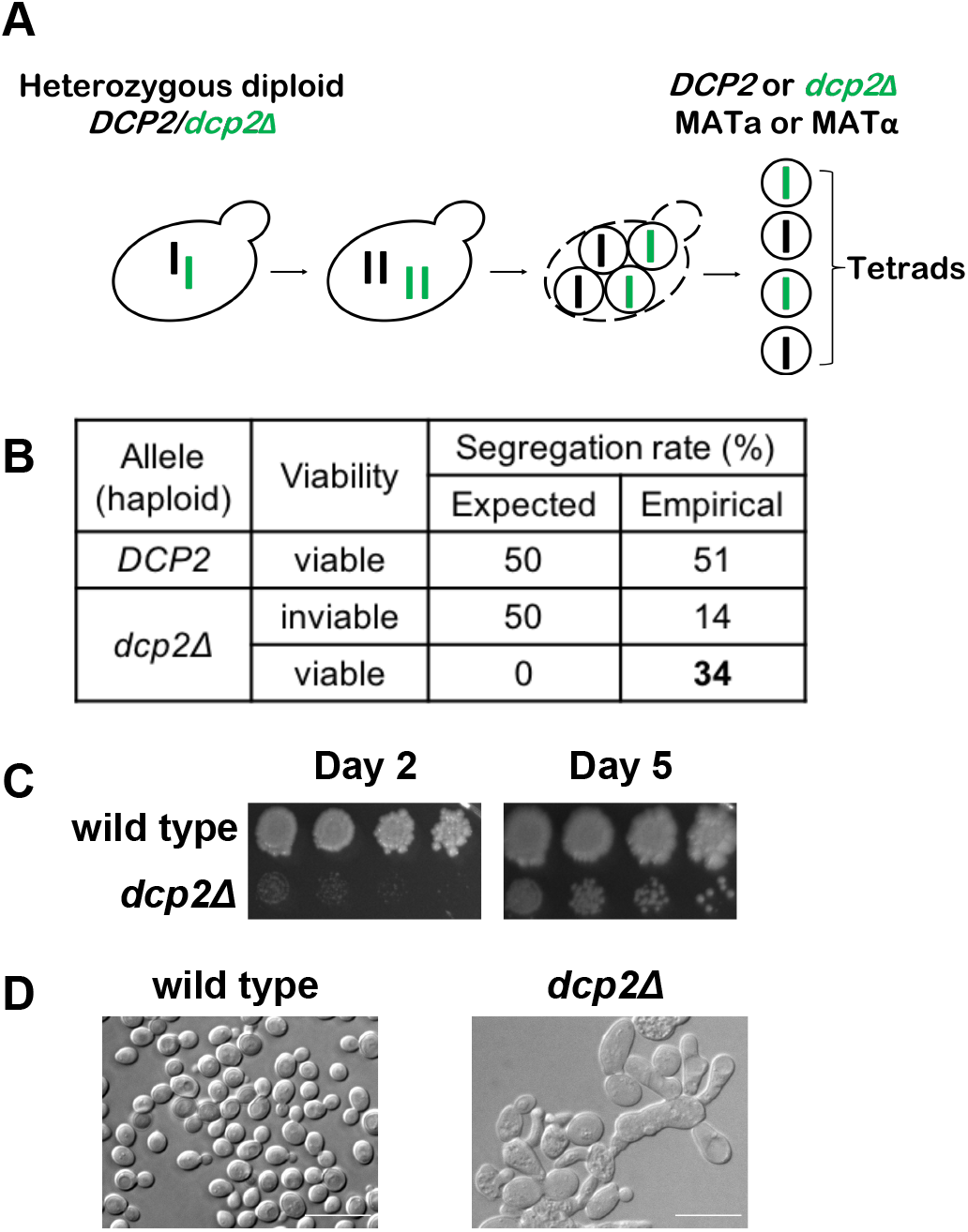
Isolation of viable *dcp2*Δ cells with severe growth and morphological defects. **(A)** Diagram of tetrad analysis of heterozygous diploid *DCP2/dcp2*Δ strain. **(B)** Tetrad dissection results in wild-type and *dcp2*Δ colonies. **(C)** *dcp2*Δ colonies resulting from tetrad dissection grow slowly. Serially diluted wild-type and viable *dcp2*Δ colonies from (B) were spotted on YPD solid media and grown at 30°C for the indicated times. **(D)** *dcp2*Δ cells resulting from tetrad dissection have morphological defects. Cells were grown at 30°C until OD_600_ reaches 0.3-0.4. Samples were diluted in YPD and examined by light microscopy. Bar represents 10μm.

To further examine the growth and morphology of the recovered *dcp2*Δ strains, they were serially diluted, spotted on YPD, and cultured at 30°C. Although viable, the d*cp2*Δ strains grow extremely slowly compared to wild type (Figure 1C). Examination through light microscopy revealed irregular and heterogeneous cell morphology, with many elongated cells in clumps (Figure 1D). Additionally, multiple vacuole-like organelles of different sizes accumulated in these cells. Taken together, this suggests that *DCP2* is required for normal growth and morphology of budding yeast (see discussion).

### Experimental evolution of *dcp2*Δ strains results in improved growth and morphology

To understand the function of *DCP2* that affects cell growth, we decided to identify suppressors of the growth defect of *dcp2*Δ. We used an experimental evolution approach that allows cells to accumulate mutations and enriches for suppressors that are advantageous for fitness in the absence of *DCP2*. For this, we used four haploid progeny, each derived from a different tetrad (meiosis). Each haploid *dcp2*Δ strain was used to start duplicate liquid cultures. Once these cultures reached saturation, we diluted them into new media for several iterations (Figure 2A). Throughout the experimental evolution process, growth of all *dcp2*Δ populations was examined both by spotting serially diluted cultures on solid media (data not shown) and by measuring OD_600_ of cells growing in liquid media (Supplemental Figure 1). During the course of the experimental evolution, we observed growth improvement at the 90^th^ generation, and further growth improvement was observed at the 180^th^ generation (Supplemental Figure 1). However, in most cases, the growth improvement from 180 generations to 270 generations was minimal (Supplemental Figure 1). Thus, we stopped the experimental evolution process after approximately 270 generations and further analyzed these *dcp2*Δ populations (evolved *dcp2*Δ). All eight evolved *dcp2*Δ populations grew better than their parental non-evolved *dcp2*Δ strain, although not as well as the wild-type strain (Figure 2B). The doubling time of the eight evolved *dcp2*Δ populations is 1.5 to 2-fold longer than that for the wild-type strain, but much shorter than the doubling time of four non-evolved *dcp2*Δ strains, which could not be calculated in the 16-hour period of the experiment due to the extreme slow growth. Similar to the growth improvement, the morphological defects in non-evolved *dcp2*Δ strains are partially restored in evolved *dcp2*Δ populations (Figure 2C and Supplemental Figure 2). Evolved *dcp2*Δ cells had a more homogenous morphology, were less elongated, and less clumped compared to non-evolved *dcp2*Δ cells. These results suggest that that the experimental evolution of *dcp2*Δ successfully selects suppressor mutations that confer the growth improvement of *dcp2*Δ strains.

**Figure 2.**
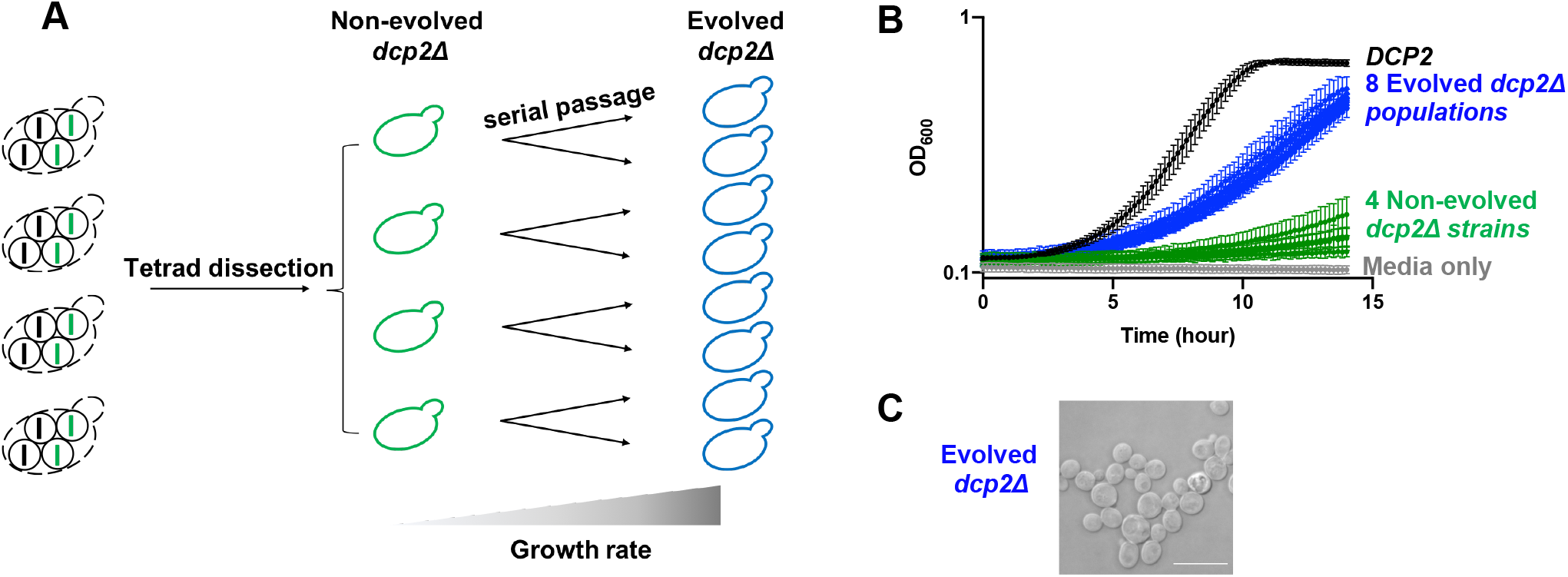
Experimental evolution of *dcp2*Δ strains results in growth and morphological improvement. **(A)** Diagram of experimental evolution. Four *dcp2*Δ strains (middle) isolated from distinct tetrads (left) were subject to serial passage. Two replicate samples of *dcp2*Δ strains were transferred to new media in iterations until the growth rate increases (right). **(B)** The *dcp2*Δ growth defect is partially improved in evolved *dcp2*Δ populations (blue) compared to non-evolved *dcp2*Δ strains (green). Cells were grown at 30°C and OD_600_ was measured every 10 minutes for ~15 hours. Shown is the average OD_600_ from replicate cultures and their standard deviations, plotted on a log scale. n=8 for *DCP2*, n=4 for each non-evolved *dcp2*Δ, and n=2 for each evolved *dcp2*Δ **(C)** Morphological defects are partially restored in evolved *dcp2*Δ populations. A representative microscopic image of an evolved *dcp2*Δ population is shown. Bar represents 10μm.

### Whole genome sequence analysis identifies suppressors of *dcp2*Δ lethality

To identify suppressor mutations that confer growth improvement to *dcp2*Δ we performed whole genome sequence (WGS) analysis on evolved *dcp2*Δ strains. We suspected that the evolved populations were genetically heterogeneous, which complicates the analysis and interpretation of WGS. Thus, for each evolved *dcp2*Δ population, we picked a single colony to generate eight genetically homogeneous evolved *dcp2*Δ isolates. As we observed in the evolved *dcp2*Δ populations, all eight evolved *dcp2*Δ isolates grew better than their non-evolved *dcp2*Δ counterparts (Figure 3A). We then sequenced the genomes of the eight evolved *dcp2*Δ isolates as well as the starting diploid *DCP2/dcp2*Δ strain (average genome coverage 112-fold), and identified mutations that were present in the evolved isolates, but not in the starting diploid (Figure 3B and Supplemental Figure 3A). Each evolved *dcp2*Δ isolate contains nonsynonymous mutations in two to six genes that are not present in the heterozygous diploid *DCP2/dcp2*Δ strain. All of them were point mutations, including substitution and deletion/insertion of a small number of bases. We did not detect any larger deletions or aneuploidy (Supplemental Figure 3B) that is often accompanied by the deletion of an essential gene [20].

**Figure 3.**
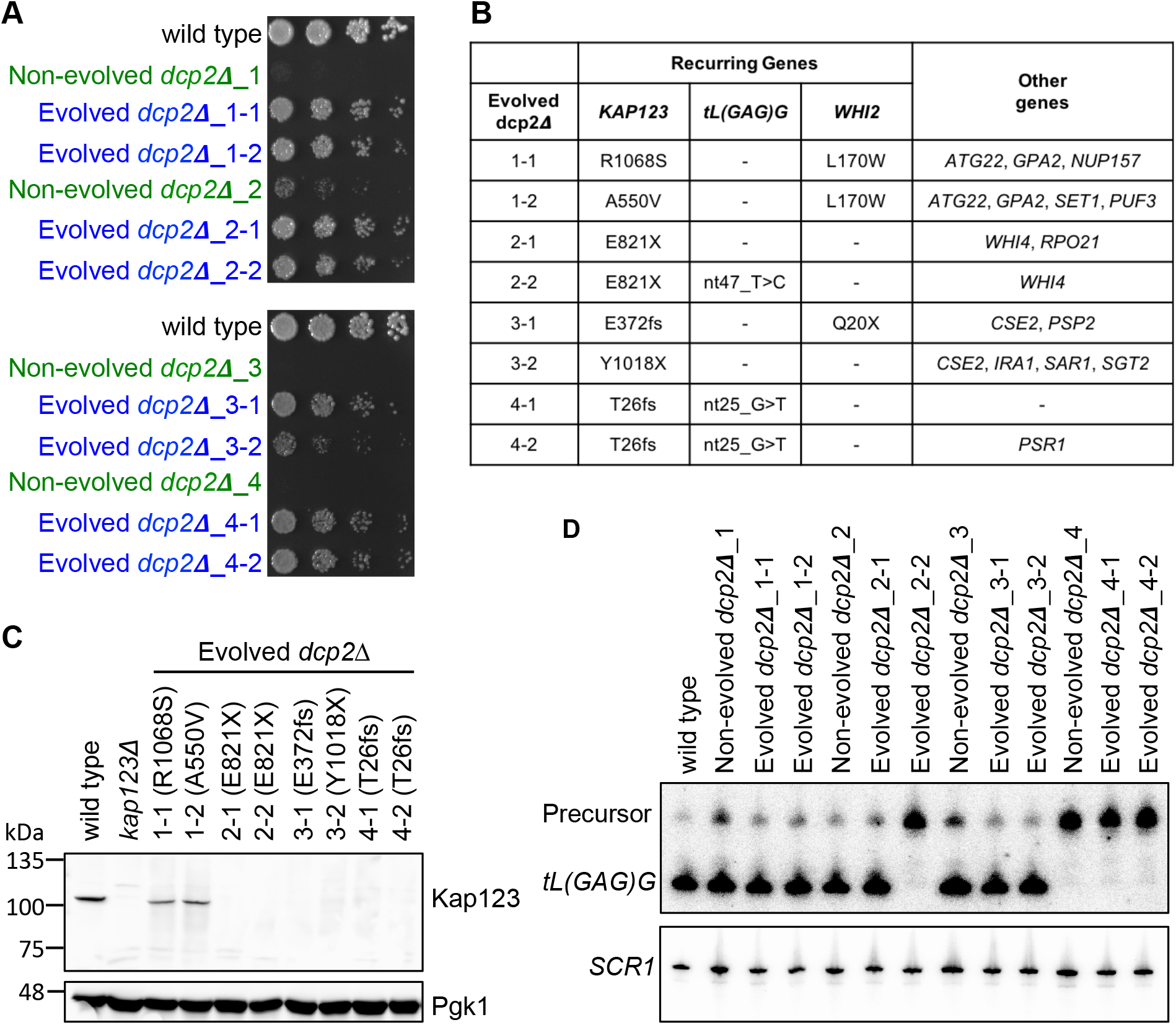
Whole genome sequencing identifies multiple null mutations in *KAP123* and *tL(GAG)G*. **(A)** The growth defect is partially improved in evolved *dcp2*Δ isolates. A single colony was isolated from each evolved population (blue) and haploid starting strain (green). Each of these genetically homogeneous strains was serially diluted, spotted on YPD solid media, and grown at 30°C. Shown is the growth at day 2. **(B)** Whole genome sequences were determined for the eight evolved *dcp2*Δ isolates and compared to the *DCP2*/*dcp2*Δ starting diploid. Nonsynonymous mutations that are not present in the starting diploid are listed. **(C)** Six of the evolved isolates have null mutations in *KAP123* that generate a premature stop codon and they do not express Kap123. A representative western blot analyzing the expression level of Kap123 (top) from the indicated strains is shown. Pgk1 (bottom) is used as loading control. **(D)** Three of the evolved isolates have null mutations in of *tL(GAG)G* and do not express mature *tL(GAG)G* tRNA. A representative northern blot of *tL(GAG)G* tRNA (top) from the indicated strains is shown. *SCR1* is used for loading control (bottom).

Strikingly, all eight evolved *dcp2*Δ isolates had a mutation in the *KAP123* gene. In total, six different *kap123* alleles were identified from the evolved *dcp2*Δ isolates. Isolates 2-1 and 2-2 are derived from the same haploid spore and both contained the *kap123-E821X* nonsense mutation. We conclude that this mutation arose very early, before the duplicate cultures were started. Similarly, each pair of evolved isolates shared at least one mutation, which must have arisen early, but also differed from its sister isolate by additional mutations (see discussion).

Four *kap123* mutant alleles have either a nonsense mutation or a frame-shift mutation that generates a premature stop codon, and thus are likely loss of function mutations. Two of the mutations are missense mutations, A550V and R1068S, that both affect conserved residues that are structurally important (Supplemental Figure 4A). Western blot analysis showed that the Kap123 protein was not detectable from the six evolved *dcp2*Δ isolates harboring nonsense or frame-shift mutation, implying destabilization of mRNA, protein, or both. In contrast, Kap123-A550V and Kap123-R1068S were expressed (Figure 3C).

**Figure 4.**
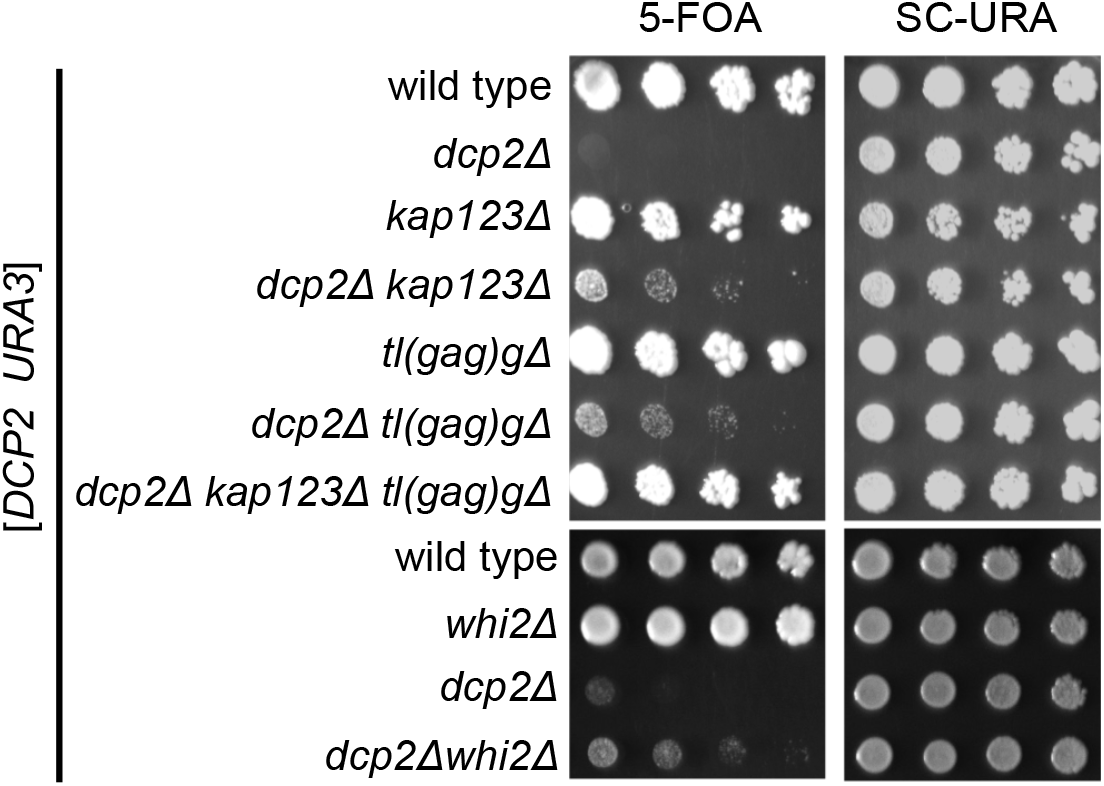
Null mutations of *KAP123*, *tL(GAG)G* or *WHI2* are sufficient to suppress the lethality of *dcp2*Δ. Growth assay showing that the lethality of *dcp2*Δ is suppressed by *kap123Δ, tl(gag)g*Δ or *whi2*Δ. Null mutations of *KAP123* and *tL(GAG)G* have an additive effect on the growth. Each strain was made by sporulation of a double heterozygous diploid transformed with a *URA3* plasmid expressing *DCP2*. Haploid progeny were serially diluted and spotted on 5FOA and SC-URA (control) solid media. Shown is the growth of a representative haploid strain.

In addition to *KAP123*, we identified multiple alleles of *tL(GAG)G* and *WHI2* in our evolved isolates. One of the *whi2* alleles introduces an early stop codon (*whi2-Q20X*) and thus is likely a loss of function allele. The *tL(GAG)G* gene encodes a leucine tRNA with a GAG anticodon, and both of the mutations are predicted to disrupt tRNA folding (Supplemental Figure 4B). Northern blot analysis indicated that the mutant tRNA was not expressed (Figure 3D). In addition, the mutant pre-tRNA appeared more abundant, suggesting that the structural perturbations in *tL(GAG)G* may interfere with 5’ end processing by RNase P and/or 3’ end processing RNase Z.

Overall, these data suggest that each of the evolved *dcp2*Δ isolates contain loss of function mutations in *KAP123*, *tL(GAG)G*, and/or *WHI2*, as well as other mutations of unclear significance.

### Null mutations of *KAP123*, *tL(GAG)G*, or *WHI2* are sufficient for viability of *dcp2*Δ

For genes that were mutated in multiple evolved *dcp2*Δ isolates, we tested whether a null mutation of each gene is individually sufficient to suppress *dcp2*Δ lethality. Because our whole genome sequencing indicated that *dcp2*Δ strains quickly accumulated suppressors, we were careful to minimize selection for undesired spontaneous suppressors that would complicate the analysis of the desired potential suppressors (*kap123*Δ, *tl(gag)g*Δ and *whi2*Δ). We therefore started with the heterozygous *DCP2*/*dcp2*Δ diploid strain that we had sequenced, and which contained wild-type *KAP123*, *WHI2*, and *tL(GAG)G* genes. We then knocked out one of the alleles of *KAP123*, *WHI2*, or *tL(GAG)G* to generate *DCP2/dcp2Δ KAP123/kap123Δ, DCP2/dcp2Δ WHI2/whi2*Δ, and *DCP2/dcp2Δ tL(GAG)G/tl(gag)g*Δ diploids. Each of these three double heterozygous strains was then transformed with a plasmid that carried functional *DCP2* and *URA3* genes, and haploid progeny were generated and genotyped. Finally the strains were grown on media lacking uracil (selecting for the *DCP2 URA3* plasmid) or media containing 5FOA (counter-selecting against the *DCP2 URA3* plasmid). Growth on 5FOA-containing media in this assay indicates that the strain is viable in the absence of the *DCP2* plasmid. This elaborate experimental setup allowed us to determine growth without inadvertently preselecting for suppressor mutations.

As expected progeny with a functional *DCP2* gene derived from each of the double heterozygotes readily formed colonies on 5FOA (Figure 4) indicating that they could grow after losing the *DCP2 URA3* plasmid. In contrast the *dcp2*Δ progeny derived from each of the double heterozygotes failed to form colonies on 5FOA, suggesting that *DCP2* is essential for normal growth. Importantly, *dcp2Δ kap123*Δ, *dcp2Δ whi2*Δ, and *dcp2Δ tl(gag)g*Δ strains each formed small colonies on 5FOA containing media, indicating that they were able to grow after losing the *DCP2* plasmid. Thus, each of these null alleles is sufficient to suppress the *dcp2*Δ lethality.

Multiple studies have reported inadvertent mutations of *WHI2* in various yeast knockout strains [21–23]. In contrast, we could not find any other studies identifying *kap123* or *tl(gag)g* as suppressors. Thus, the effect of *whi2*Δ on *dcp2*Δ appears to be less specific than the other two suppressors. Therefore, we decided to focus on *KAP123* and *tL(GAG)G* for the further analyses.

To test whether *kap123*Δ and *tl(gag)*Δ suppressed *dcp2*Δ lethality through a common pathway we generated *dcp2Δ kap123Δ tl(gag)*Δ triple mutant with the *DCP2 URA3* plasmid. Importantly, the triple mutant grew better on 5FOA plates than either double mutants (Figure 4), indicating that *kap123*Δ and *tl(gag)g*Δ act independent of each other to improve *dcp2*Δ growth. This observation that multiple suppressors further improve growth also explains why the experimental evolution resulted in strains with multiple mutations.

### Other viable *dcp2*Δ or *dcp1*Δ strains contain similar suppressor mutations

Our WGS data suggest that *DCP2* is required for viability of *S. cerevisiae*, but this conclusion is not consistent with some previous studies reporting that *DCP2* is dispensable for survival. Some studies have attributed these differences to the use of different strains. *e.g*. it has been reported that *dcp2*Δ is viable in the W303 strain but lethal in S288C [16, 24]. However, the available genome sequences of W303 and S288C revealed that neither contains obvious null mutations in *KAP123*, *WHI2*, or *tL(GAG)G*. We therefore re-analyzed the strains used in three different studies.

First, we analyzed published RNA-seq data of a *dcp2*Δ strain in the S288C background to reveal mutations shown as mismatches between the RNA reads and wild-type genome [15]. The *dcp2*Δ strain used in that study was obtained from our lab and is derived from the non-evolved strain *dcp2*Δ_4 used in our studies. Our WGS above indicated that non-evolved strain *dcp2*Δ_4 strain had an early arising *tl(gag)g-G25U* allele, and as expected, we detected this allele in the RNA-seq reads. In addition, we detected a *kap123-T766fs* allele that is different from mutations what we identified in WGS analysis. Thus, this strain acquired the *tl(gag)g* mutation at some point before we shared it, and acquired an additional *kap123* mutation at some point before the RNA-seq was performed.

We also re-analyzed publicly available RNA-seq data for the previously reported viable *dcp2*Δ and *dcp1*Δ mutants in the W303 strain background [25]. Importantly, all of the reads from the *dcp2*Δ strain that mapped to codon 786 of *KAP123* indicated that this codon was a UAA codon instead of a UAC codon (Supplemental Figure 5A). We conclude that the *dcp2*Δ strain used in this RNA-seq experiment is a *dcp2Δ kap123-Y687X* double mutant. Similarly, in the *dcp1*Δ strain, we detected a 21 nt deletion in *KAP123* (*kap123-Δ510-516;* Supplemental Figure 5B). Moreover, the RNA reads from the control W303 strain indicated that it contained a wild-type *KAP123* gene. Thus the *kap123* mutations in the *dcp1*Δ and *dcp2*Δ strains are not inherent differences between W303 and S288C, but instead mutations that inadvertently arose in these strains.

**Figure 5.**
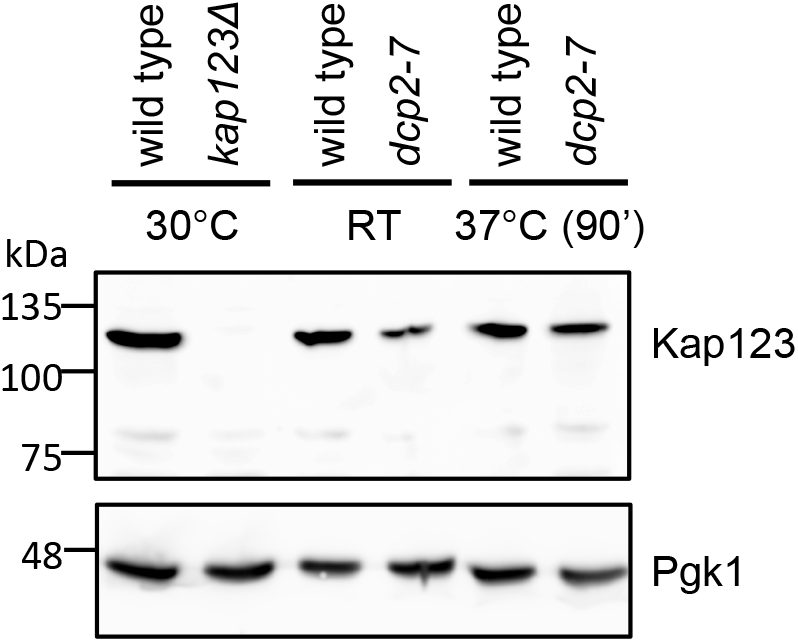
Decapping is not required to maintain Kap123 at a low nontoxic level. Expression of Kap123 is not increased in decapping deficient cells. A representative western blot analyzing the expression of Kap123 (top) is shown. Pgk1 (bottom) is used a loading control. Lane 1-2: cells were grown at 30°C, lane 3-4: cells were grown at room temperature (RT), and lane 5-6: cells were grown at RT until OD_600_ reached 0.6-0.8 then transferred to 37°C for 90 minutes.

Lastly, much of the initial investigations of decapping were carried out in a yRP strain background, and *dcp2*Δ was described as viable in this background [5, 26]. We therefore PCR amplified and sequenced the *KAP123* gene from this *dcp2*Δ strain (yRP1346) and detected that the PCR product was a mixture of *kap123-G727X* and *KAP123*. Thus, a *kap123* mutation appeared to have arisen after the *dcp2*Δ strain was created.

Overall, these results indicate that many of the previously reported viable *dcp2*Δ strains contain previously undetected mutations in *KAP123* that likely contribute to their viability. We conclude that *DCP2* is an essential gene in *S. cerevisiae* (see discussion).

### Effect of decapping defects on *KAP123* expression

While most genes in *S. cerevisiae* can be overexpressed, some “dosage-sensitive” genes cause growth defects upon overexpression. *KAP123* is among the most “dosage-sensitive” genes [27]. Thus, we considered the possibility that decapping is critical to maintain the *KAP123* mRNA and Kap123 protein at a low non-toxic level. A *dcp2*Δ strain would thus be lethal because Kap123 is overexpressed to toxic levels and a *kap123* mutation would restore viability. We tested this possibility using two different approaches. Because all of our *dcp2*Δ strains contain *kap123* mutations, we used a *dcp2-7* temperature sensitive strain, which has a wild-type *KAP123* gene (see methods) and performed western blot analysis. This showed that Kap123 was not overexpressed upon Dcp2 inactivation (Figure 5). We also checked the *KAP123* levels in the RNA-seq data from *dcp2-7* and *dcp1*Δ strains [25, 28]. The *dcp2-7* strain contains a wild-type *KAP123* allele, and although the *dcp1*Δ contains a *kap123* mutation, this in frame deletion is not expected to affect *KAP123* mRNA stability. Consistent with our western blot, neither decapping mutant strain had an increased *KAP123* mRNA level (Supplemental Figure 6A). These results indicate that suppression of the essential function of Dcp2 is not to reduce *KAP123* expression below a toxic level.

**Figure 6.**
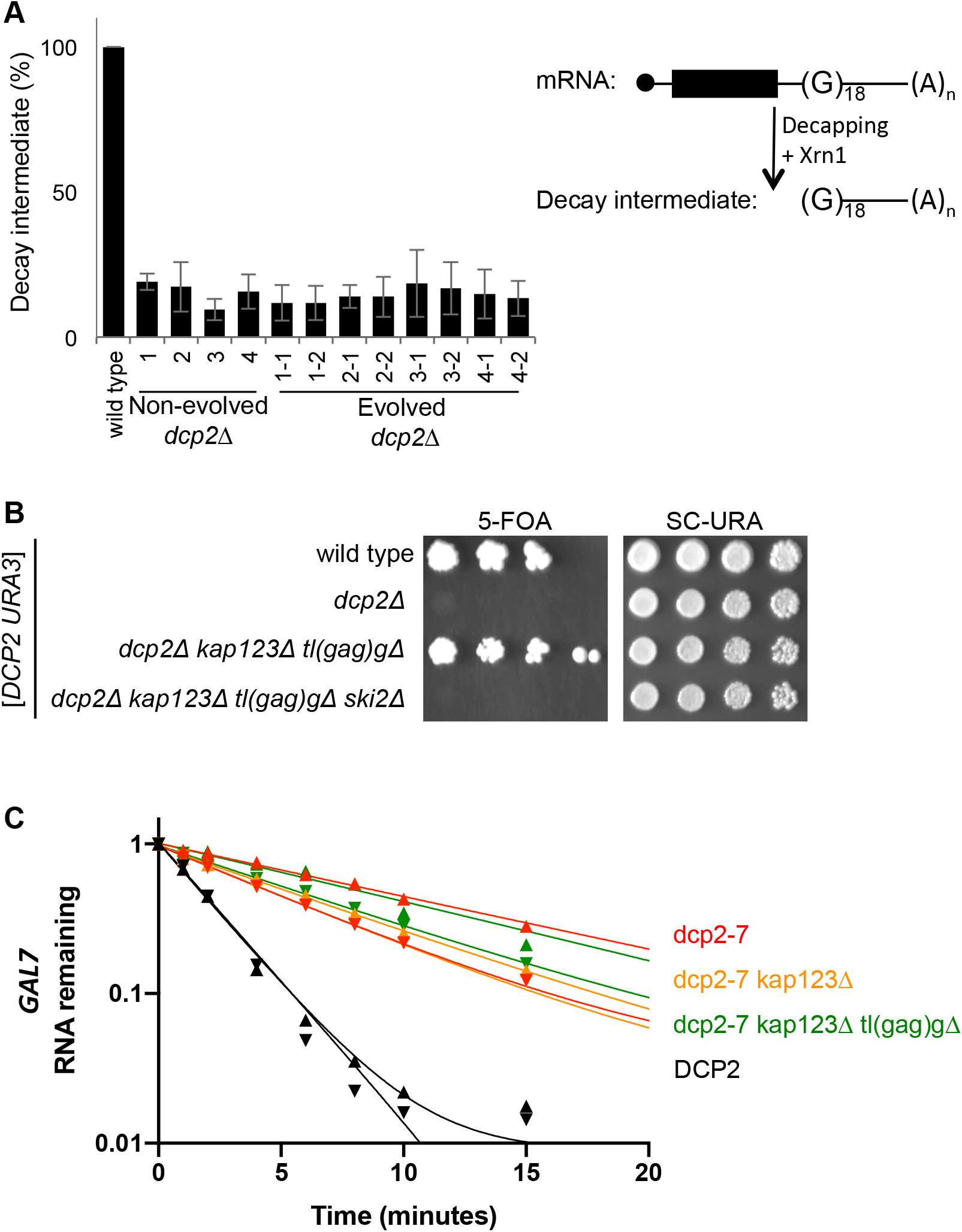
Suppressor mutations of *dcp2*Δ do not restore mRNA decay. **(A)** Decapping is defective in evolved *dcp2*Δ strains. Each strain was transformed with an *MFA2pG* reporter (right). Decapping, or endonucleolytic cleavage, followed by 5′ exonucleolytic decay of *MFA2pG* produces a decay intermediate that is a sensitive measure of decapping. Total RNA was isolated, and *MFA2pG* mRNA and decay intermediate levels were analyzed by northern blotting. Plotted is the mean ratio of decay intermediates to mRNA from three replicates and its standard deviation normalized to wild type (left). **(B)** RNA exosome-mediated decay is required in *dcp2*Δ suppressor mutants. *dcp2Δ kap123Δ tl(gag)g*Δ is no longer viable in the absence of *SKI2*. Each strain was made by sporulation of a quadruple heterozygous diploid transformed with a *URA3* plasmid expressing *DCP2*. Haploid progeny were serially diluted and spotted on 5FOA and SC-URA (control) solid media. Shown is the growth of a representative haploid strain at day 5. **(C)** The stability of *GAL7* mRNA is unaffected by *kap123*Δ and/or *tl(gag)g*Δ in *DCP2* deficient cells. Cells exponentially growing at 21°C in galactose were transferred to 37°C for 1 hour. Transcription of *GAL7* was repressed by the addition of dextrose at time 0, and cells were harvested at multiple time points. Total RNA was isolated and *GAL7* mRNA levels were analyzed by northern blotting. Plotted is remaining *GAL7* mRNA levels relative to *SCR1* levels of two biological replicates. Data point triangles are pointing up for one replicate, and down for the other.

### Suppressor mutations of *dcp2*Δ do not restore mRNA decay

We next used several assays to determine whether the suppressor mutations affected mRNA decay rates or pathways. First, we examined whether decapping activity is restored in the evolved *dcp2*Δ isolates. Several other enzymes carry out similar activities including Dcs1, Dxo1, Rai1, and the L-A viral GAG protein. Specifically, Dcs1 cleaves the cap structure that remains after the mRNA is degraded in the 3′ to 5′ direction, Dxo1 and Rai1 digest aberrant caps, and GAG transfers the cap from cellular mRNAs to viral RNA [7, 8, 29-33]. Although none of these enzymes have canonical activities for bulk cytoplasmic decapping, it appeared possible that our suppressor mutations redirected their activity to bulk cytoplasmic mRNA. To test this possibility, we compared the *in vivo* decapping activity among wild-type, non-evolved, and evolved *dcp2*Δ isolates. In this assay, we used an mRNA with a G-quadruplex structure in the 3′ UTR. This structure impedes Xrn1, and therefore any decapping results in the accumulation of a decay intermediate that is easily detectable by northern blotting ([34]). Northern blots revealed that both the non-evolved and evolved *dcp2*Δ strains lacked the decay intermediate to a similar level (Figure 6A). This result demonstrates that the evolved *dcp2*Δ strains are still defective in mRNA decapping, at least of this widely used model mRNA.

In addition to exonucleases, some endonucleases have been shown to contribute to cytoplasmic mRNA decay in yeast. For example, Ire1 and tRNA splicing endonuclease (TSEN) cleave specific mRNAs [35, 36]. One possibility is that our suppressors increase activity the activate or decrease the specificity of some endonuclease. We also entertained the idea that *tl(gag)g* mutations could cause stalling of translating ribosomes, which might trigger “no-go” cleavage by Cue2 [37, 38]. Regardless of the endonuclease involved, endonucleolytic cleavage of the mRNA upstream of the G-quadruplex initiates Xrn1-mediated decay and leads to the same decay intermediate as decapping [36–39]. Therefore, the absence of this decay intermediate indicates that the suppressors do not increase endonucleolytic decay of *MFA2pG* mRNA.

Second, it has previously been shown that if the 5′ to 3′ decapping pathway is impaired, the RNA exosome and its associated helicase Ski2 become rate limiting for mRNA decay [40]. This is reflected in growth phenotypes. Specifically, *ski2*Δ does not cause an obvious growth defect if Dcp2 is fully functional, but is lethal if decapping activity is impaired [40]. We therefore combined *ski2*Δ with *dcp2Δ kap123*Δ and *tl(gag)g*Δ. As before the *dcp2Δ kap123Δ tl(gag)g*Δ triple mutant is viable, but the *dcp2Δ kap123Δ tl(gag)gΔ ski2*Δ quadruple mutant is inviable (Figure 6B), indicating that cytoplasmic RNA exosome activity is required even in the presence of *dcp2*Δ suppressors. This suggests that the RNA exosome is the major mRNA degrading activity in the *dcp2Δ kap123Δ tl(gag)g*Δ mutant. Overall, the *MFA2pG* assay and the *ski2* synthetic lethality indicate that *kap123*Δ and *tl(gag)*Δ do not result in hyperactivation of a novel mRNA degradation pathway.

Third, we tested the effect of *kap123*Δ and *tl(gag)*Δ on stability of a specific mRNA. Because all our *dcp2*Δ strains already have suppressor mutations, we again used the temperature-sensitive *dcp2-7* allele in this experiment and compared the stability of the *GAL7*, *GAL1* and *GAL10* mRNAs in a *dcp2-7* strain to their stability in *dcp2-7 kap123*Δ and *dcp2-7 kap123Δ tl(gag)g*Δ strains. For this experiment, the expression of the *GAL* mRNAs was induced by growth in the presence of galactose, the decapping enzyme was then inactivated by incubating the cultures at 37°C. Finally, glucose was added to inhibit transcription from the *GAL* genes, and RNA was isolated in a time course. This revealed that the *GAL* mRNAs were each degraded more slowly in the *dcp2-7* strain compared to a *DCP2* control as expected [5, 40]. However, when comparing the *dcp2-7 kap123*Δ and *dcp2-7 kap123Δ tl(gag)g*Δ mutants to *dcp2-7* we detected no differences (Figure 6C and supplemental Figure 7). In both of the suppressed strains the mRNA half-lives were similar to the *dcp2-7* single mutant, and more stable than the wild-type control.

**Figure 7.**
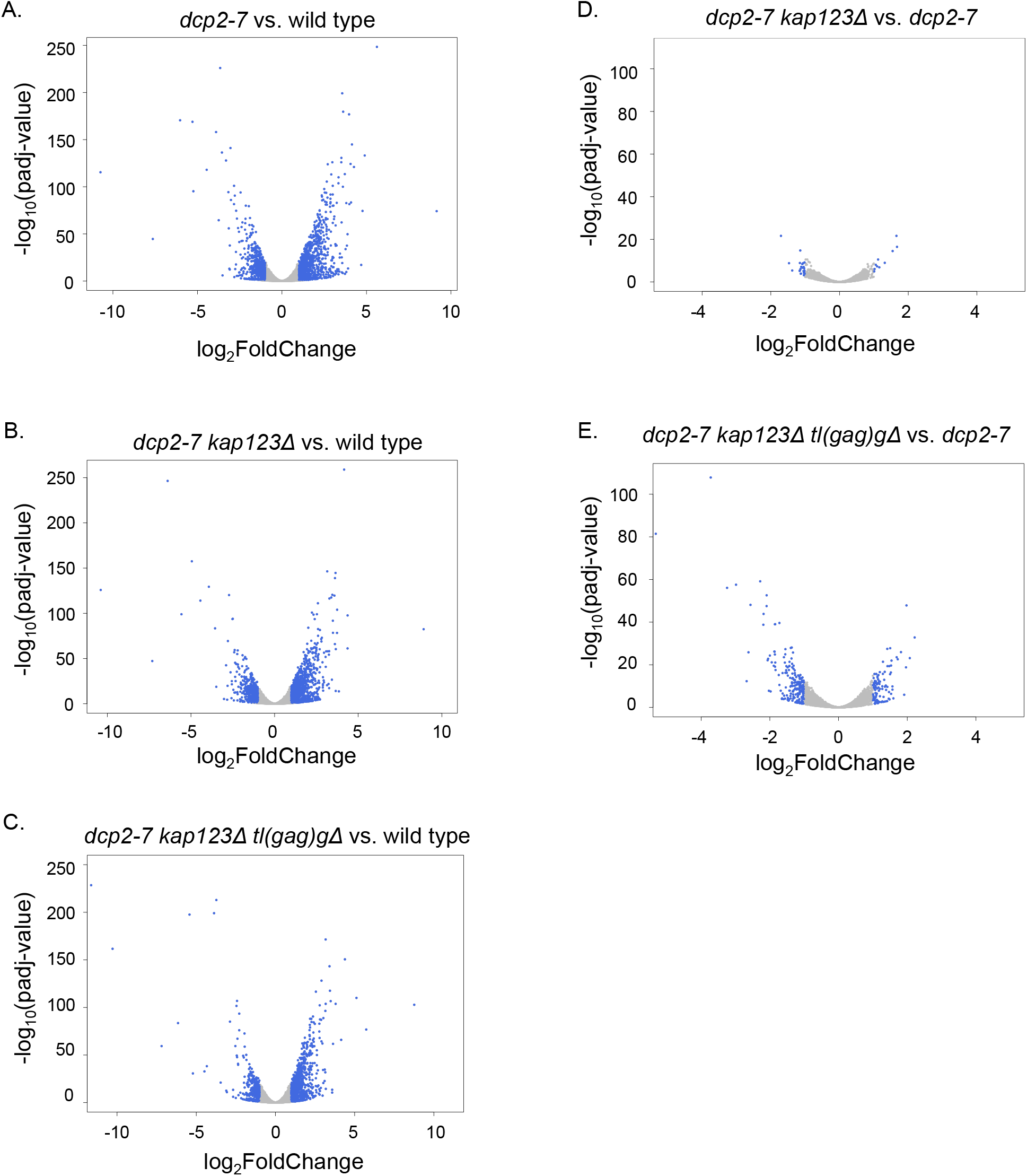
Transcriptome analysis of *dcp2* mutants. **(A-C)** Volcano plots showing differential expression of 7127 annotated genes in *dcp2-7* mutants versus wild type. **(D-E)** Volcano plots showing differential expression of 7127 annotated genes in *dcp2-7* suppressor mutants versus *dcp2-7* strain. (A-E) Transcripts that significantly changed at least 2-fold (adjusted p-value<0.05) are in blue. Note the difference in scales between panels A to C and D and E.

Taken together, these results show that unlike previously isolated suppressors [5, 17–19] our suppressor mutations do not restore decapping or mRNA decay rates. Instead, while *kap123*Δ and *tl(gag)g*Δ affect the viability of *dcp2*Δ, they do not detectably affect cytoplasmic mRNA decay.

### Suppressors ameliorate mRNA and ncRNA defects in a *dcp2* mutant

To gain a better understanding of how *dcp2*Δ suppressors affect the transcriptome we performed global gene expression analysis. Biological triplicates of wild type, *dcp2-7*, *dcp2-7 kap123*Δ, and *dcp2-7 kap123Δ tl(gag)g*Δ were subjected to RNA-seq analysis after enrichment of poly(A)^+^ RNAs. As previously reported, *dcp2-7* causes widespread disruption of the transcriptome [28]. In our analysis the *dcp2-7* strain shows 1004 annotated genes significantly (adjusted p-value < 0.05) upregulated by ≥ 2-fold and 618 annotated genes downregulated by ≥ 2-fold (Figure 7A). As in previous analysis, similar numbers of genes were up- and down-regulated, which we interpret to indicate that the effects are dominated by indirect effects, instead of reflecting altered stability. A similar but slightly lower number of genes were affected in the *dcp2-7 kap123*Δ mutant where 972 annotated genes were upregulated by ≥ 2-fold and 550 annotated genes were downregulated ≥ 2-fold compared to wild type (Figure 7B). Even fewer significant changes in the global transcripts levels were detected in the *dcp2-7 kap123Δ tl(gag)g*Δ mutant where 752 annotated genes were upregulated by ≥ 2-fold and only 340 annotated genes were downregulated by ≥ 2-fold (Figure 7C). However, even the triple mutant showed widespread defects. We interpret these results to indicate that the suppressors have significant but modest effects on the transcriptome.

To determine whether specific genes were affected by *kap123*Δ and *tl(gag)g*Δ, we compared the gene expression profile of *dcp2-7 kap123*Δ and *dcp2-7 kap123Δ tl(gag)g*Δ to the *dcp2-7* strain. Unlike the comparison to wild type, there are only a few genes differentially expressed. In *dcp2-7 kap123*Δ mutant, there were only 10 annotated genes upregulated by ≥ 2-fold and 20 annotated genes downregulated by ≥ 2-fold (Figure 7E) when compared to the single mutant *dcp2-7*. In the *dcp2-7 kap123Δ tl(gag)g*Δ mutant, 102 annotated genes were upregulated by ≥ 2-fold and 201 annotated genes downregulated by ≥ 2-fold compared to *dcp2-7* (Figure 7E). Thus, suppression of growth defects is accomplished by surprisingly modest effects on the transcriptome.

To determine whether the suppression mechanism engages the regulation of certain biological processes or cellular functions, we examined if any specific biological processes are enriched in transcripts whose expression significantly changed at least 2-fold (adjusted p-value < 0.05) via Gene Ontology (GO) enrichment analysis. When the gene expression of the *dcp2-7*, *dcp2-7 kap123*Δ, and *dcp2-7 kap123Δ tl(gag)g*Δ mutants were compared to wild type, the top five most enriched GO terms for upregulated transcripts were related to RNA and ribosomal RNA modifications, while the top five most enriched GO terms in downregulated transcripts were related to cell cycle processes (Supplemental Figure 8). Because the GO analysis results were similar among all three *dcp2* mutants, we looked into transcripts that are differentially expressed between d*cp2-7* and *dcp2-7* with suppressor mutations. The GO term analysis on transcripts upregulated in *dcp2-7 kap123*Δ relative to *dcp2-7* revealed enrichment of the biological process related to cell adhesion, while the same analysis on transcripts downregulated revealed the enrichment of the biological process related to organization of nucleic acids (Supplemental Figure 9). Interestingly, the only GO category enriched in transcripts downregulated in *dcp2-7 kap123Δ tl(gag)g*Δ relative to *dcp2-7* is rRNA modification (Supplemental Figure 9). This was particularly striking, because this GO category was upregulated in *dcp2-7* and although it was still upregulated in *dcp2-7 kap123Δ tl(gag)g*Δ, the upregulation was significantly less. This GO category therefore correlated with the growth effect. The genes in this GO category that were affected were mostly small nucleolar RNA (snoRNA) encoding genes. Thus, snoRNA genes are significantly disregulated in *dcp2-7* compared to wild type and this dysregulation is partially alleviated by suppressor mutations.

**Figure 8.**
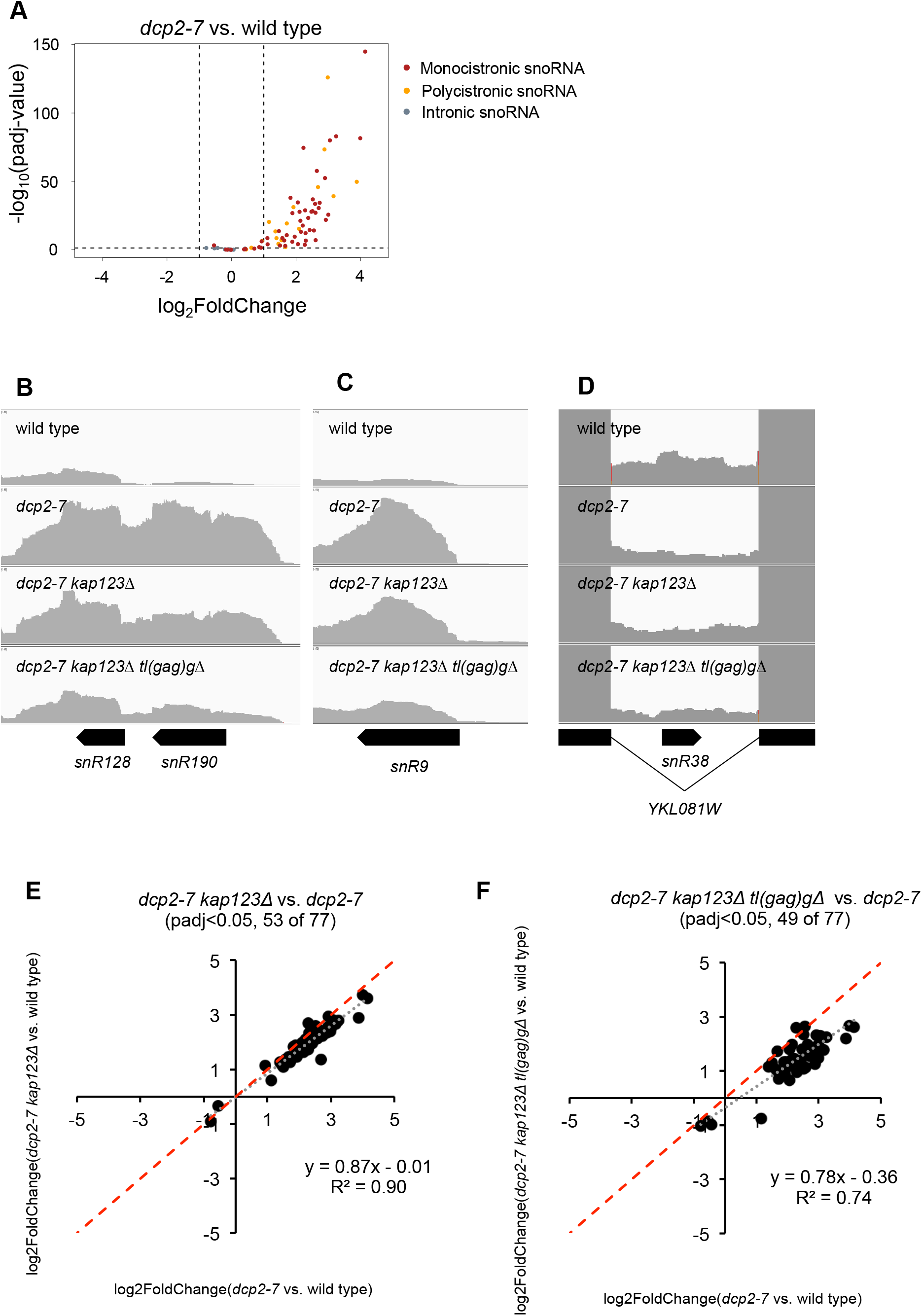
Suppressors ameliorate expression defects from snoRNA loci in *dcp2-7* mutant. **(A)** The expression of mono- and polycistronic snoRNA loci are upregulated in *dcp2-7* mutant. A volcano plot showing differential expression of snoRNA genes in *dcp2-7* strain versus wild type. Monocistronic snoRNA are in red, polycistronic snoRNA genes in orange and intronic snoRNA genes in gray. Dotted lines represent cut-off values for differential gene expression, log_2_(Foldchange) ≥ 1 and adjusted p-value < 0.05. **(B-D)** Upregulation of mono- and polycistronic snoRNA loci in *dcp2-7* mutant were reduced by *kap123*Δ and further by *kap123Δ tl(gag)g*Δ. RNA-seq read coverage for the snoRNA gene loci *snR128*-*snR190* (dicistronic), *snR9* (monocistronic), and *snR38* (intronic) are shown. **(E-F)** Defective snoRNA expression in *dcp2-7* mutant was alleviated by *kap123*Δ and further by *kap123Δ tl(gag)g*Δ. Plotted is the differential expression of snoRNA loci in suppressor mutants versus *dcp2-7* mutant (relative to wild type). Of 77 snoRNA genes in *S. cerevisiae*, significantly changed (adjusted p-value <0.05) 53 and 49 transcripts were plotted, respectively. The red dotted line with slope 1 is the predicted outcome if *kap123*Δ and *tl(gag)g*Δ had no effect. The grey dotted line with slope <1 depicts linear regression analysis results.

**Figure 9.**
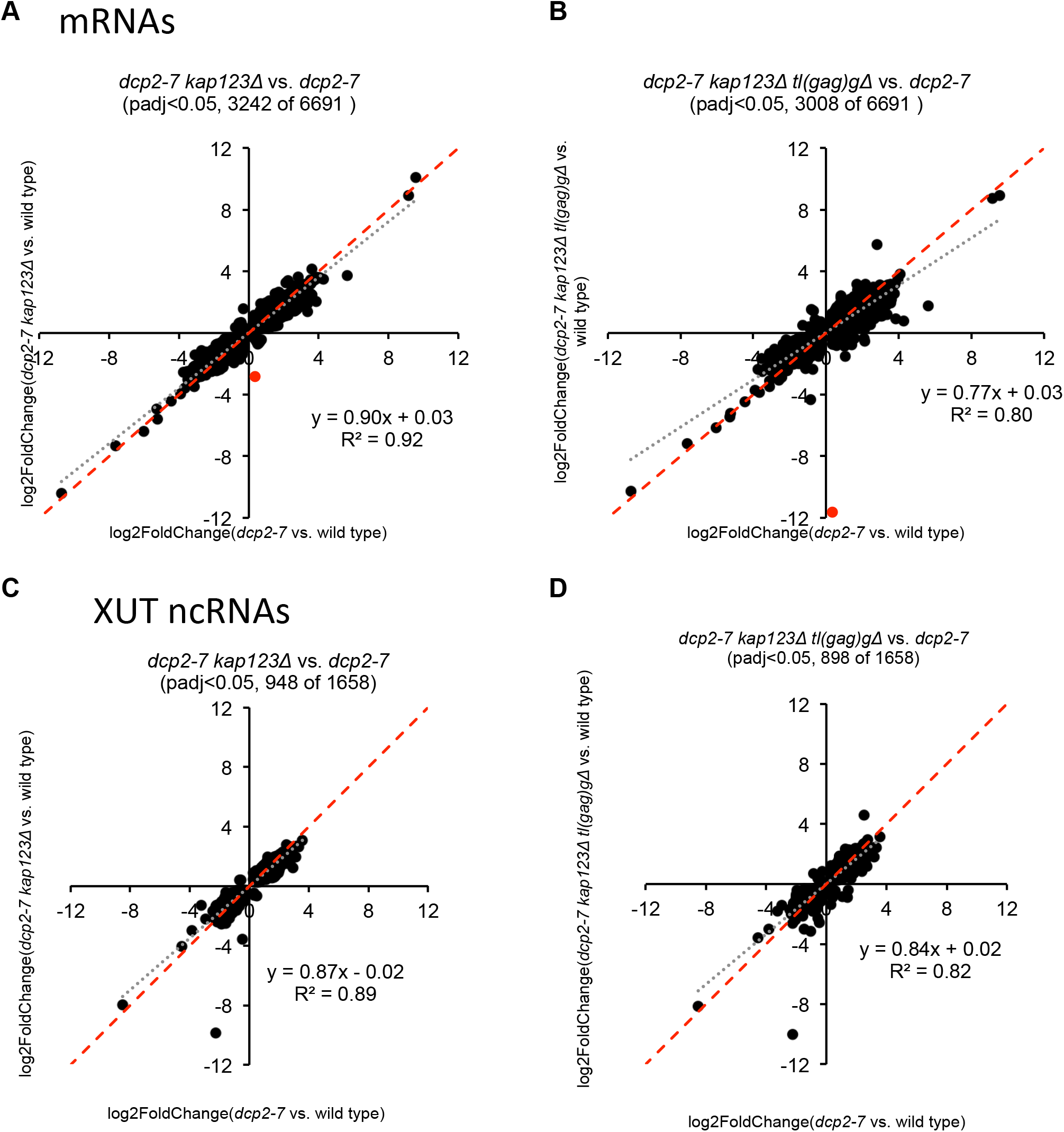
Suppressors ameliorate defects in expression of protein coding mRNA and XUT non-coding RNA in *dcp2-7* mutant. **(A-B)** Defective mRNA expression in *dcp2-7* mutant was alleviated by *kap123*Δ and further by *kap123Δ tl(gag)g*Δ. Plotted is the differential expression of mRNA loci in suppressor mutants versus *dcp2-7* mutant (relative to wild type). Of 6691 mRNA genes in *S. cerevisiae*, significantly changed (adjusted p-value < 0.05) 3242 and 3308 transcripts were plotted, respectively. Red dot indicates differential expression of *KAP123* **(C-D)** Defective XUT expression in *dcp2-7* mutant was alleviated by *kap123*Δ and further by *kap123Δ tl(gag)g*Δ. Plotted is the differential expression of XUT loci in suppressor mutants versus *dcp2-7* mutant relative to wild type. Of 1658 XUT genes in *S. cerevisiae*, significantly changed (adjusted p-value <0.05) 948 and 898 transcripts were plotted, respectively.

snoRNAs can be classified by function and conserved elements into C/D box snoRNAs that direct RNA methylation and H/ACA snoRNAs that direct pseudouridylation of RNA, and both classes of snoRNAs appeared to be affected. snoRNAs can also be divided by gene organization into monocistronic, polycistronic, and intron-encoded snoRNAs. Each of these categories is processed, but Dcp2 is not thought to play a role in snoRNA processing. Instead, the monocistronic snoRNAs are transcribed independently as ^7m^G capped precursors. These pre-snoRNAs are then 5′ matured into ^2,2,7m^G-capped snoRNAs by further methylations [41] or into a 5′ phosphorylated snoRNA by cleavage by the endonuclease Rnt1 and the exonuclease Rat1 [42, 43]. Similarly, polycistronic precursors are separated into individual snoRNAs by Rnt1, and then 5′ processed by Rat1[44, 45]. Finally intron-encoded snoRNAs are processed from the spliced out and debranched intron [46]. Both polycistronic (Figure 8A and B) and monocistronic (Figure 8A and C) snoRNA loci were upregulated in *dcp2-7*, but most intron-encoded snoRNA loci were not (Figure 8A and D). Moreover, inspection of read coverage at snoRNA loci indicated that what we detected was an upregulation of transcripts from snoRNA loci that were 5′ and/or 3′ extended, rather than the mature snoRNAs (Figure 8B and C). Importantly, the loci for both uncapped (Figure 8B) and capped (Figure 8C) mature snoRNAs were affected. These observations indicate that the defect does not reflect a role of Dcp2 in snoRNA processing.

To quantitate the effect of the suppressors, we plotted the log2(fold change) of *dcp2-7* versus that in *dcp2-7 kap123*Δ or *dcp2-7 kap123Δ tl(gag)g*Δ. If the *kap123*Δ and *tl(gag)g*Δ had no effect, this should result in data points along a line with a slope of 1. On the other hand, complete restoration of the *dcp2-7* defect would result in a line with a slope of 0. Linear regression of the data revealed slopes of 0.87 and 0.78 with high correlation coefficients (R^2^=0.90 and R^2^=0.74) indicated that the snoRNA transcripts were decreased 13% and 22% in the double and triple mutant relative to the *dcp2-7* single mutant (Figure 8E and F). These results showed that defective snoRNA regulation in *dcp2* mutant cells is ameliorated by suppressors.

Although snoRNAs were enriched among genes whose expression was increased in *dcp2-7* and partially restored in *dcp2-7 kap123Δ tl(gag)g*Δ, we sought to determine whether this effect was restricted to snoRNAs. With this goal in mind we repeated the analysis of log2(fold change) in *dcp2-7* versus the suppressed strains for annotated genes, which are mostly protein coding genes (Figure 9A and B), and XUTs (Figure 9C and D). XUTs are ncRNAs transcribed by RNA polymerase II and increased in abundance upon *xrn1*Δ [47]. Because XUTs are transcribed by RNA polymerase II, they should be capped, and thus have to be decapped before they can become Xrn1 substrates. This analysis revealed that the disruption of mRNA and XUT expression detected in *dcp2-7* is also partially restored in the *dcp2-7 kap123*Δ strain, and even more in the *dcp2-7 kap123Δ tl(gag)g*Δ strain. Overall, our data showed global transcriptomic changes in the Dcp2-deficient strain and that this defect is mitigated by suppressors. Moreover, Dcp2 affects expression of a large proportion of the transcriptome, including both mRNAs and non-coding RNAs such as snoRNAs and XUTs.

## DISCUSSION

### Dcp2 is essential for continuous growth

Cytoplasmic mRNA turnover is an important process that regulates gene expression, and Dcp2 carries out a key step. Previous studies have described *DCP2* as either an essential gene, or a non-essential gene, and in some cases the difference has been attributed to a genetic difference between the two widely used yeast strains, S288C and W303 [5, 14–16]. Our studies indicate that *DCP2* is indeed essential, but that suppressor mutations in several different genes can allow for viability. We further show that previous studies have inadvertently used *dcp1*Δ or *dcp2*Δ strains that contain spontaneous *kap123* suppressors. However, similar analysis from other *dcp2* alleles (e.g. *dcp2-7* and *dcp2-N245* [28, 48]) and from strains with mutations in decapping regulators (*lsm1*Δ and *pat1*Δ etc. [48]) indicated that they contained a wild-type *KAP123* gene. This suggests that only very severe decapping defects select for *kap123* mutations.

The starting *DCP2/dcp2*Δ heterozygote used in our tetrad analysis does not contain these suppressors, but a large fraction of the resulting haploid colonies do. It appears implausible that ungerminated and metabolically inactive spores accumulate suppressor mutations to high frequency, without accumulating a large number of other random mutations. Therefore, we conclude that *dcp2*Δ spores must be capable of germinating and dividing for several generations. Strong selective pressure during early generations leads to the selection for suppressors that then allow further growth into visible colonies. It takes approximately 20 generations to form a colony, allowing sufficient time for some initial suppressors to occur [49]. The need for an initial suppressor for a spore to form a visible colony also explains why approximately 30% of *dcp2*Δ spores did not form visible colonies. Thus, Dcp2 is likely not absolutely essential for spore germination and extremely slow growth, but in practical terms it is essential for continuous culture under lab conditions, since *dcp2*Δ cultures cannot be maintained without suppressors.

### Experimental evolution identifies novel bypass suppressors of *dcp2*Δ

Our experimental evolution approach to identifying suppressors in the decapping enzyme is distinct from previous genetics screens. All previous screens isolated suppressors of a partial defect in the decapping enzyme (*dcp2-7* or *dcp1-2*) and these suppressor mutations restored mRNA decapping by improving the function of the partially defective Dcp1/Dcp2 enzyme [5, 17–19]. Thus, these previous screens were helpful in identifying regulators of Dcp1/Dcp2 but they did not provide much insight into the importance of decapping for growth and cellular homeostasis. We designed our experimental evolution approach to complement these previous studies, reasoning that the use of a complete deletion of the *DCP2* gene would preclude selecting suppressors that restored Dcp1/Dcp2 function. Experimental evolution further allows for selection of suppressors with small effects on growth. Small effects on growth may not result in a readily observable effect on colony size in a traditional screen, but are selected slowly but surely in experimental evolution.

We identified nonsynonymous mutations in 16 different genes. Some of these mutations are likely to just have occurred randomly, and may not affect *dcp2*Δ growth. However, three genes were mutated multiple times and we concentrated our further analysis on those. We show that *kap123*Δ, *tl(gag)g*Δ and *whi2*Δ loss of function mutations are each sufficient to suppress the lethality of *dcp2*Δ. One advantage of experimental evolution is that multiple mutations can arise that act additively or even synergistically to further improve growth. This seems to have occurred since six of the eight isolates contain mutations in both *KAP123* and either *tL(GAG)G* or *WHI2*. Furthermore, we confirmed that the *dcp2Δ kap123Δ tl(gag)g*Δ triple mutant grew better than either the *dcp2Δ kap123* or *dcp2Δ tl(gag)g*Δ double mutant. Another advantage of experimental evolution is that it allows multiple mutations to arise where mutation of a particular gene can only improve growth after some other gene is mutated. We did not find clear evidence for this, although this might be the case for some of the mutations we only found once. Although we cannot completely trace the order in which each mutation arose, the comparison of duplicate evolved isolates derived from a common haploid spore indicates some mutations arose early and others arose later (Supplemental Figure 10A). This indicates that in lineage 4 a *tl(gag)g* mutation arose before a *kap123* mutation, while in lineage 2 the genes were mutated in the reverse order.

Although we have not formally proven it, several lines of evidence suggest that some of the 13 genes mutated only once also affect *dcp2*Δ viability. First, evolved isolate 2-1 contains a *kap123* mutation, but wild-type *WHI2* and *tL(GAG)G* genes, yet grows just as well as some of the isolates with two suppressors. Thus, isolate 2-1 likely contains an additional suppressor. The only nonsynonymous mutation that differs between isolate 2-1 and the unevolved parent is a single amino acid change in RNA polymerase II (*rpo21-R1281C*), and thus this may be a suppressor. Second, in lineage 3, both the *kap123* and *whi2* mutations arose late. This suggests that some other early mutation might have allowed us to recover the non-evolved starting strain. The only nonsynonymous mutation shared between isolates 3-1 and 3-2 is a frame-shift in *CSE2*, which encodes a subunit of the mediator complex. Third, isolate 4-2 contains a mutation in the start codon for Psr1, which forms a complex with Whi2. Further analyses will be required to definitively determine whether these 13 other mutations also affect *dcp2*Δ growth.

### Bypass suppressors of *dcp2*Δ do not restore decapping or mRNA decay rate to normal

Previous genetic screens of suppressors of *dcp1* or *dcp2* defects have identified “enhancer of decapping” and “inhibitor of decapping” genes that restored decapping to a Dcp1/Dcp2 complex with point mutations [5, 17–19]. We expected to either isolate mutations that activate alternative mRNA degradation pathways or mutations that allow survival despite slow mRNA decay. Follow up experiments indicate that we isolated the latter. We observed that the suppressor mutations isolated here do not appear to affect the pathway of mRNA decay, nor the rate. Instead they appear to improve viability while maintaining the relatively slow RNA exosome-mediated 3′ to 5′ decay of mRNA.

### Bypass suppressors of *dcp2*Δ partially restore the transcriptome

Finally, we studied the effect of *kap123*Δ and *tl(gag)g*Δ on the transcriptome. Limitations of this study include that we had to use a *dcp2-7* strain because *dcp2*Δ strains could not be cultured without suppressors arising and thus could not be used as a single mutant control to compare to the double mutants. Although *dcp2-7* causes a decapping defect at the restrictive temperature, this defect is not complete. Comparing *dcp2-7* to wild type, we observed global disruption in transcriptome homeostasis, confirming previous publications [15, 16, 25]. We suspect that the detected changes are dominated by indirect effects and thus cannot be used to identify which transcripts are differentially decapped. Strikingly, GO analysis indicated a significant enrichment of snoRNA loci among the transcripts that were restored by *kap123*Δ and *tl(gag)g*Δ. snoRNA processing pathways are well-established and do not involve decapping of snoRNA transcripts. In fact, some mature snoRNAs retain the cap structure while for other snoRNAs the Rnt1 endonuclease removes the cap. Both of these classes of snoRNAs were affected. For example, the snR9 snoRNA is well-established to retain its cap [50] and is not processed by Rnt1 [42] while both snR190 and snR128 are 5′ processed by Rnt1 [44], yet all three are similarly affected by Dcp2 inactivation (Figure 8B and C). Thus, Dcp2 inactivation appears to indirectly affect transcripts from the snoRNA loci, and the *kap123*Δ and *tl(gag)g*Δ suppressors partially suppress this defect. Although snoRNAs were enriched among transcripts whose partial restoration upon *kap123Δ tl(gag)g*Δ reaches statistical significance, we observed by linear regression that mRNAs and XUTs follow a similar trend, but perhaps are not as sensitive to *kap123Δ tl(gag)g*Δ and thus individually do not reach significance. Our findings that suppressor mutations ameliorate the effect of decapping defects, and that most previous transcriptome-wide studies inadvertently used strains with suppressors imply that most previous studies likely underestimated the transcriptome-wide effects of decapping defects.

## MATERIALS AND METHODS

### Strains, plasmids, and oligonucleotides

The *DCP2/dcp2*Δ heterozygous diploid in the BY4743 (S288C) background was obtained from Research Genetics and all other strains (Supplemental Table 1) used are derived from it through standard genetic procedures. Plasmids were generated by standard procedures and are listed in Supplemental Table 2. Oligonucleotides (Sigma-Aldrich) used in this study are listed in Supplemental Table 3.

### Yeast growth conditions

Yeast was grown either in standard yeast extract peptone (YEP) media containing 2% dextrose or galactose or in synthetic complete media lacking amino acids (Sunrise Science) as required. G418 (0.67 mg/ml), hygromycin B (325 U/mL), or clonNAT (100 μg/ml) was added to YEP+dextrose media to select for knockouts. Cells were incubated at 30°C unless otherwise indicated. The *dcp2-7* cultures were incubated for 60 or 90 minutes at 37°C to inactivate the decapping enzyme for the *GAL* mRNA stability and transcriptome sequencing experiments, respectively.

To induce sporulation, diploid cells were grown in nutrient-depleted media for 4-5 days. Sporulated cells were resuspended in water with Glusulase (Perkin Elmer). This reaction was incubated at 30°C for 30 minutes. Ascus digestion was terminated using water. Haploids were obtained either by tetrad dissection or by random spore isolation [51].

Experimental evolution was initiated from four haploid *dcp2*Δ strains, each derived from a different tetrad. Duplicate 5 ml cultures of each of the four haploid *dcp2*Δ strains were inoculated in YEP containing 2% dextrose, G418 (167 mg/L), and ampicillin (50 mg/L). Cultures were grown at 30°C until the OD_600_ of the culture reached > 8.5. 10 μl of this culture (containing on the order of 10^6^ yeast cells) were then transferred into 5 mL of fresh media of the same type. This culturing and 500-fold dilution was repeated 30 times. A 500-fold (or 2^8.97^) dilution represents approximately 9 doublings. Thus, after 30 cycles of culture and dilution the cultures had gone through approximately 270 generations.

For the growth assay on solid media, exponentially growing cells were serially diluted (5-fold) and spotted on the indicated media. For the growth assay in liquid media, exponentially growing cells were diluted to OD_600_ of 0.1, and transferred to a 96-well plate. Cells were incubated at 30°C in a BioTek’s Synergy™ Mx Microplate Reader. OD_600_ was measured every 10 minutes for ~15 hours. Collected data were processed through Gen5 (BioTek).

### Microscopy

To examine cell morphology, exponentially growing cells were analyzed on an Olympus BX60 microscope.

### Whole genome sequencing analysis

Total genomic DNA was isolated from exponentially growing cells using a phenol-chloroform extraction method and further purified with the use of a DNeasy Blood & Tissue Kit (Qiagen) and a MasterPure™ Yeast DNA Purification Kit (Lucigen). PE150 libraries of the evolved strains were prepared and sequenced by Novogen.

To identify mutations, sequencing reads were trimmed with Trim Sequences (http://hannonlab.cshl.edu/fastx_toolkit/), quality checked with FastQC (http://www.bioinformatics.babraham.ac.uk/projects/fastqc/), and mapped with Bowtie2 [52] to *S. cerevisiae* reference genome R64-1-1(www.emsembl.org). The overall alignment rate was ~91 to 99%. Before calling variants, BAM datasets for individual *dcp2*Δ strain and heterozygous diploid *DCP2*/*dcp2*Δ strain were merged using MergeSamFiles (http://broadinstitute.github.io/picard/). Datasets were further processed for left realignment through BamLeftAlign (https://arxiv.org/abs/1207.3907). In order to call all the variants, FreeBayes (https://arxiv.org/abs/1207.3907) for detection and SnpEff 4.3 [53] for annotation were used. Integrated Genome Viewer (https://software.broadinstitute.org/software/igv/download) was used to inspect candidate SNPs. True mutations were differentiated from sequencing errors and pre-existing SNPs by being supported by the consensus of the reads in the evolved isolate(s), but not by the reads from the other evolved isolates or the heterozygous diploid *DCP2*/*dcp2*Δ strain that we had previously sequenced. MiModD Deletion Calling (https://sourceforge.net/projects/mimodd/) was used to search for deletions, which identified the *ura3Δ, his3Δ, met15Δ, lys2*Δ, and *leu2*Δ deletions of BY4743, but no other deletions. MiModD Coverage Statistics (https://sourceforge.net/projects/mimodd/) was used to measure coverage depth by chromosome to search for aneuploidy.

### Protein analysis

Total protein was extracted in IP50 buffer (50mM Tris-HCl (pH7.5), 50mM NaCl, 2mM MgCl_2_, 0.1% Triton X-100) with 0.007% beta-mercaptoethanol and 0.00174% PMSF, and complete protease EDTA-free mini tablet (Roche) by bead beating and analyzed by western blot. Blots were probed with anti-Kap123 at 1:5,000 [54, 55] and anti-Pgk1 at 1:10,000 (Invitrogen) and developed using Amersham ECL Prime (GE Healthcare). Images were acquired and analyzed using an ImageQuant™ LAS 4000 biomolecular imager (GE Healthcare) and ImageQuant TL image analysis software.

### RNA analysis using northern blotting

For analyzing the steady-state RNA level, cells exponentially growing at 30°C were harvested. For analyzing RNA stability, *dcp2-7* mutants were grown in YEP containing 2% galactose at 21°C and transferred into a 37°C incubator for 1 hour to inactivate the decapping enzyme. Cells were washed with YEP, and dextrose (40% stock solution) was added to a final concentration of 2% to repress transcription of the *GAL* genes. While cells were incubated at 37°C, samples were collected at the indicated time points and immediately frozen.

For RNA preparation, the harvested cell pellet was lysed by vortexing with glass beads. RNA was purified through two rounds of phenol/chloroform/LET and one additional chloroform extraction, and ethanol precipitated. Total RNA was analyzed through northern blotting. Briefly, 10 μg of total RNA was analyzed by electrophoresis on denaturing gels, either 1.3% agarose/formaldehyde gels for mRNA analysis or 6% polyacrylamide (19:1) 8M urea gel for tRNA analysis, as indicated. RNA was transferred to a nylon membrane and probed with P^32^ 5′ end labeled oligonucleotides. Blots were imaged by phosphorimaging on a Typhoon FLA 7000 (GE Healthcare), and quantitated using imagequant software.

### Transcriptome analysis

For RNA sequencing analysis, cultures (biological triplicates) of exponentially growing cells were transferred into a 37°C incubator and incubated for 90 minutes to inactivate the decapping enzyme before harvesting. Total RNA was extracted using the hot phenol method [56]. For sequencing, poly(A)^+^ selected RNA was used to construct a library for PE150 Illumina sequencing. Reads were submitted to SRA under project PRJNA626686. Raw reads were trimmed using Trim Galore! (http://www.bioinformatics.babraham.ac.uk/projects/trim_galore/), quality checked with FastQC (http://www.bioinformatics.babraham.ac.uk/projects/fastqc/), and mapped to reference genome R64-1-1 with TopHat2 [57]. Gene expression level was determined with featureCounts [58] for genes identified in the annotation file from Ensembl (www.emsembl.org) and for XUTs (http://vm-gb.curie.fr/XUT/index.htm) [47]. Differential gene expression was determined through DESeq2 [59]. Gene ontology analysis was performed through the GO Term Finder (Version 0.86).

### Finding *kap123* mutations in published RNA-seq data

To determine whether previously published RNA-seq experiments inadvertently used *kap123* mutant strains and to identify the mutations, we downloaded raw RNA-seq reads from EBI (https://www.ebi.ac.uk/ena). Reads were trimmed with Trim Galore!, quality checked with FastQC, and then aligned with TopHat2 to the R64-1-1 reference genome. Aligned reads were analyzed in IGV. This identified the *kap123-Y687X* in three data sets from a *dcp2*Δ W303 strain (SRR4163304, SRR4163305, SRR4163306). For the *dcp1*Δ datasets (SRR4163301, SRR4163302), the TopHat alignment suggested a small deletion, but failed to precisely identify it. Aligning the same datasets with Bowtie2 in very sensitive local mode did precisely identify the deletion as a 21bp deletion mediated by a GCGGAACC repeat in the wild-type gene. We found no mutations in the *dcp1*Δ or *dcp2*Δ strains for any of the other genes that are mutated in our evolved isolates. Similarly, analyzing the *dcp2*Δ RNA-seq data from reference [15] (SRR364981) we identified the same *tl(gag)g* mutation as in our evolved isolates 4-1 and 4-2 and a novel *kap123* mutation. We did not find *kap123* mutations in wild-type controls (SRR4163289, SRR4163290, SRR4163291), *xrn1*Δ (SRR4163307, SRR4163308, SRR4163309) *dcp2-7* (SRR2045250, SRR2045251, our RNA-seq data), *dhh1*Δ (SRR6362787), *pat1*Δ (SRR6362781), *lsm1*Δ (SRR6362784), *dcp2-N245* (SRR6362793), *dcp2-N245-E153Q* (SRR6362796), *dcp2-N245-E198Q* (SRR6362799), *scd6*Δ (SRR7162931), *caf1*Δ (SRR7174202), or *dhh1*Δ (SRR3493892, SRR4418659) [25, 28, 48, 60-63]. This suggests that only very severe decapping defects select for *kap123* suppressors.

## ACKNOWLEDGEMENTS

This work was supported by NIH R01 grant GM099790 to AvH. We thank members of the van Hoof laboratory for critical discussions, Kate Travis for performing pilot experiments for the experimental evolution, and Jaeil Han for insightful comments throughout. Antibodies to Kap123 were a kind gift from Valerie Doye (Institut Jaque Monod) and initially developed by Michael Rexach (University of California, Santa Cruz). The yRP1346 *dcp2*Δ strain was a kind gift from Roy Parker (University of Colorado). We thank William Margolin (UTHSC-Houston) for microscope usage, Kevin Morano (UTHSC-Houston) for microplate reader usage, and Cesar Arias, Diana Panesso and An Dihn (UTHSC-Houston) for whole genome sequencing of the *DCP2/dcp2*Δ starting diploid strain.

